# Spatiotemporal characterisation of information coding and exchange in the multiple demand network

**DOI:** 10.1101/2024.10.07.617103

**Authors:** Hamid Karimi-Rouzbahani, Anina N. Rich, Alexandra Woolgar

## Abstract

The multiple-demand network (MDN), a brain-wide system with nodes near sensory and higher-order cognitive regions, has been suggested to integrate and exchange task-related information across the brain, supporting cognitive task performance. However, the profile of information coding and the role of each node within this network in information exchange remain unclear. To address this, we combined fMRI and MEG data in a challenging stimulus-response mapping task. Using multivariate pattern analysis (MVPA), we decoded various forms of task information, including coarse and fine stimulus details, motor responses, and stimulus-response mapping rules, across the MDN and visual regions. Early in the task, visual regions responded to large physical differences in stimuli, while later on, fine stimulus information and rules were encoded across the MDN. To assess information exchange between regions, we developed Fusion-RCA, a novel connectivity analysis method based on fMRI-MEG fusion profiles. Our findings revealed significant transfer of fine stimulus information, rules, and responses, but little evidence for the transfer of coarse stimulus information. These results highlight distinct information encoding patterns within MDN nodes and suggest that the anterior cingulate cortex (ACC) plays a key role in distributing task-relevant information. This study offers new insights into the dynamic function of the MDN and introduces Fusion-RCA as a powerful tool for exploring brain-wide information transfer.

## Introduction

Humans are able to engage in intelligent goal-directed behaviour across different scenarios. To achieve this, the brain must combine information from multiple systems, from the sensory input of multiple modalities, through to memory, attention and motor processes. These systems are distributed across the brain, and a key question in understanding intelligent behaviour is how such information is integrated and coordinated.

An excellent candidate for this high-level executive function was suggested to be a network of regions known as the MDN (Duncan et al., 2020), also called the frontoparietal control network (e.g., Marek and Dosenbach, 2018), or cognitive control network/networks (e.g., Cole and Schneider, 2007). These regions include areas in the intraparietal sulcus (IPS), anterior insula/frontal operculum (AI/FO), inferior frontal sulcus (IFS), and dorsal anterior cingulate cortex (ACC), and overlap considerably with the frontoparietal resting-state network (Assem et al., 2020). They are active in many cognitive tasks (Fedorenko et al., 2013; Assem et al., 2020) and are widely thought to have a key role in supporting intelligent flexible behaviour (Jung and Haier, 2007; Duncan et al., 2010; Colom et al., 2010). For example, the extent of damage to these regions (and not nearby language regions) linearly predicts deficits in fluid intelligence (Woolgar et al., 2010; Woolgar et al., 2018). Fluid intelligence refers to the ability to think abstractly and solve novel tasks/problems and is predictive of a wide range of cognitive abilities (Baltes et al., 1999).

In line with its putative role in supporting a wide range of goal-directed behaviours, the MDN shows remarkable flexibility in adaptively coding information across a wide range of tasks (Zheng et al., 2024). Univariate functional Magnetic Resonance Imaging (fMRI) studies showed activation in this network across cognitive tasks, including those involving memory, maths, conflict detection, visual discrimination and more (Stiers et al., 2010; Fedorenko et al., 2013; Assem et al., 2020), suggesting its involvement in cognitively challenging tasks regardless of specific content (Duncan and Owen, 2000; Dosenbach et al., 2006). Multivariate fMRI studies have demonstrated that the MDN differentiates a range of different types of information (including stimuli, rules and responses) across tasks (Crittenden et al., 2016; Woolgar et al., 2016; Shashidhara et al., 2024). Moreover, within tasks, MDN responses are strongly shaped by task relevance (Zheng et al., 2024) and difficulty (Stiers et al., 2010; Woolgar et al., 2011a; Woolgar et al., 2015b). For example, representations of identical stimuli were enhanced in the MDN when attention was directed to objects (Woolgar et al., 2015b) or object features (Jackson et al., 2017; Wisniewski et al., 2023; Moerel et al., 2024). Representations across the MDN were also enhanced for hard vs. easy stimulus (Woolgar et al., 2011a; Woolgar et al., 2015b) and rule (Woolgar et al., 2015a). By combining transcranial magnetic stimulation (TMS) and fMRI, it has further been demonstrated that the right dorsolateral prefrontal cortex node (dlPFC) of the MDN plays a causal role in the dominance of task-relevant information in the system specifically by upregulating task-relevant codes (Jackson et al., 2021).

There is also evidence that connectivity and information coding in the MDN can predict behavioural performance. For example, in an auditory detection task, pre-stimulus connectivity between auditory sensory areas and cingulo-opercular nodes of the MDN (i.e., ACC and AI/FO) correlated with the accuracy of behavioural responses (Sadaghiani et al., 2015). In a stimulus-response (SR) mapping study, where different rules determined the mapping between stimuli and response buttons, the information held by the MDN correlated with behavioural responses, such that when the response was incorrect, the MDN also represented incorrect information (Woolgar et al., 2019; see also Robinson et al. 2022). These results suggest an important role for the MDN in processing task-related information and supporting behaviour.

A key proposal has been that the MDN could underpin flexible goal-directed behaviour by functionally connecting distinct and distant cognitive systems of the brain (Jung and Haier, 2007; Cole et al., 2013; Duncan et al., 2020; Zheng et al., 2024). This requires functional connections between the areas involved. Indeed, in addition to co-activation and flexible encoding of information across tasks, there is also evidence from resting-state fMRI studies that, compared to non-MD areas, the nodes of the MDN are highly connected (Cole et al., 2010; Power et al., 2013; Assem et al., 2020). They follow a small-world network structure (Bullmore and Sporns, 2009), which can facilitate access to distant parts of the brain through relatively shorter pathways compared to a regular network where information needs to pass many nodes to move across the brain. Such connections provide a potential mechanism by which the MDN could bring together different sources of information in arbitrary ways (Higo et al., 2011; Cole et al., 2013; Duncan et al., 2020).

While the above studies have shown resting-state and functional connectivity between the nodes of the MDN, empirical evidence for information exchange across these nodes is missing. Specifically, it remains unclear whether co-activation and co-encoding of information across the nodes of the MDN reflect *information exchange* between those nodes. Alternatively, it could be that the nodes of the MDN become active simultaneously and encode similar information without necessarily exchanging information. Information coding and information exchange do not necessarily co-occur (Anzellotti and Coutanche, 2018). Therefore, it is unclear whether the encoded information held within nodes of the MDN moves across the brain. Moreover, the profile of information coding across the MDN and the nature of the transferred information remain unclear. To address this, we developed a new fusion-based representational connectivity analysis (Fusion-RCA) to evaluate the exchange of information between MDN nodes. We combined the logic of Granger causality with the spatiotemporal richness of fused MEG-fMRI data to quantify the information flow between the nodes of the MDN, and between these nodes and the visual system. We used this to examine the exchange of stimulus, rule and response information in a complex stimulus-response mapping task. We also separated the representation of stimulus information into coarse (i.e., the stimuli presented on the left vs. right side of the visual field) and fine (i.e., the stimuli presented inner vs. outer of the visual field) components, which we predicted may have differential representations in the MD system based on previous work showing higher MDN coding of stimulus information that is perceptually difficult to discriminate (Woolgar et al., 2011; Woolgar et al., 2015b).

We found that different nodes of the MDN encoded task-related information with subtly different dynamics and observed information transfer from sensory regions to MDN and its circulation within the MDN and back to early visual cortex (EVC). Fine stimulus information was exchanged more strongly than coarse stimulus information across the MDN. Rule information flowed bidirectionally between MDN and the visual area while response information flowed from ACC to posterior nodes of the MDN and EVC. We found a discriminable coding and connectivity profile in the cingulo-opercular (CO) and the frontoparietal (FP) nodes of the MDN, which aligns with previous distinctions (Crittenden et al., 2016). Our results provide insights into how information is exchanged within and beyond the MDN in support of flexible rule-based behaviour. We also introduce a new brain connectivity method for studying exchange of different types of information with high spatial and temporal resolution in humans.

## Results

To characterise information coding and transfer across the MDN, we used a paradigm that allowed us to evaluate information about stimuli, rules and button-press responses (Robinson et al., 2022). Participants were presented with one stimulus and rule cue on each trial after which they were supposed to press one of four buttons depending on a previously learned stimulus-response mapping (Figure 1A). On each trial, the participant had to discriminate the location of the stimulus (one of four options, two on the left of fixation, two on the right) and apply the rule indicated by the cue colour (e.g., cue colour 1 or 2 indicated Rule 1 should be applied; colour 3 or 4 indicated Rule 2 should be applied; counterbalanced across participants; Figure 1B). Participants performed the task with relatively high performance (accuracy: mean = 80.78%, sd = 9.29%; correct reaction time: mean = 1692 ms, sd = 284 ms), indicating that they had learned the task and could successfully implement the rules, despite the task complexity. Participants failed to respond on an average of 2.6% (sd = 3.04%) of trials.

**Figure 1.**
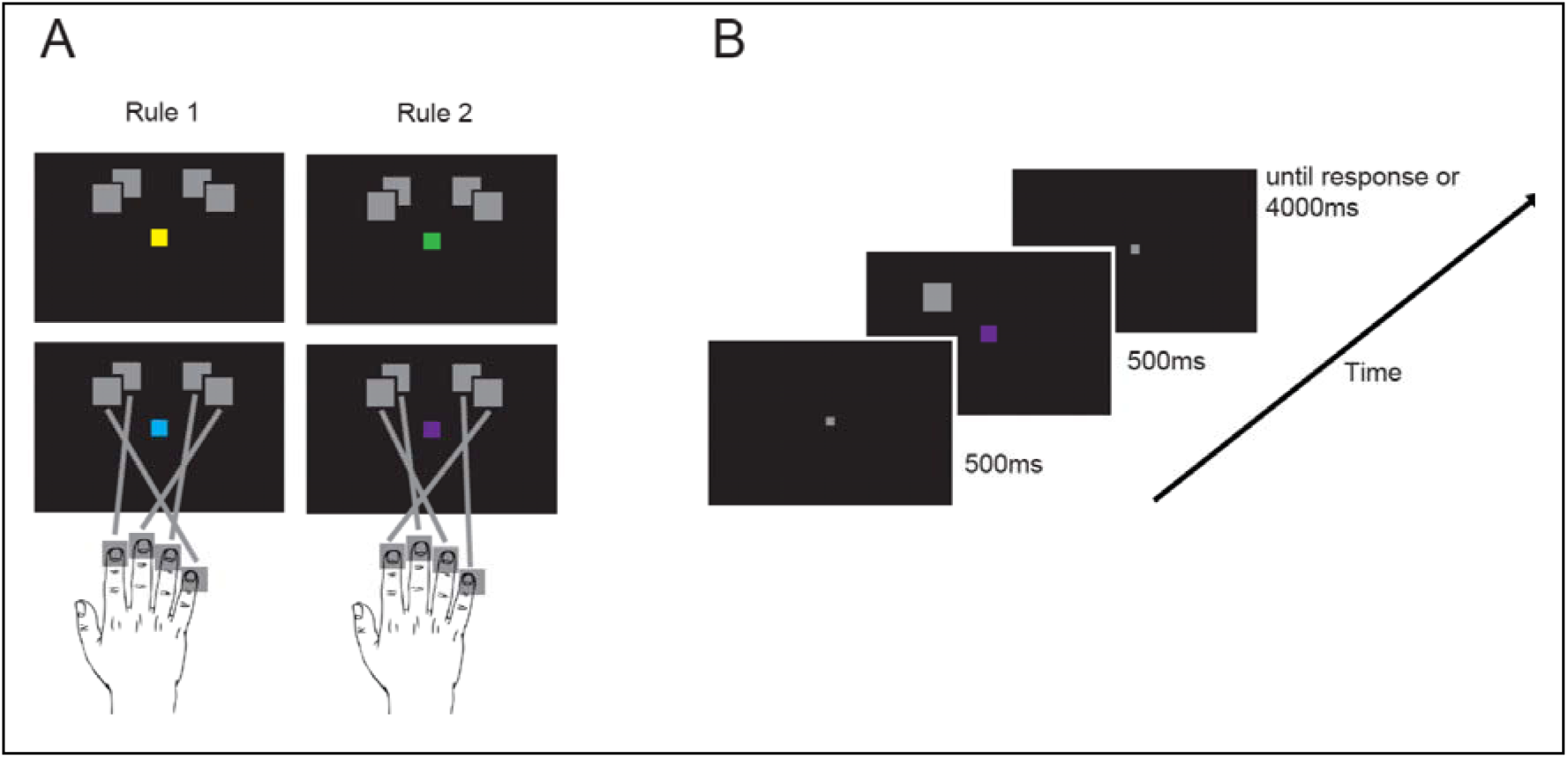
Experimental paradigm and the definition of different types of information (figure reproduced from Robinson et al., 2022). (A) Experiment was a stimulus-response mapping task, where participants had to press the button that corresponded to each stimulus. Participants were trained to perform two distinct stimulus-response mappings (i.e., rules) which were indicated by the colour of fixation square (two per rule). We defined the coarse stimulus information as the distinction between the stimuli presented on the left vs. right side and the fine stimulus information as the distinction between the stimuli presented in the inner vs. outer positions in the visual field. Rule information refers to the distinction between the two rules and response information refers to the distinction between inner vs. outer finger press responses. (B) Participants were cued about which rule to apply on a trial-by-trial basis. Each trial started with a fixation cross, followed by a stimulus-rule display and then the response screen.

### Analysis 1: multivariate pattern analysis in space and time

As an initial step, we evaluated the coding of different types of information using multivariate pattern analysis (MVPA). For this, we used Region of Interest (ROI)-specific fMRI decoding (Figure 2A) in early visual cortices, the lateral occipital complex (LOC), and the regions that form the MDN. Next, we performed time-resolved decoding on existing MEG data (Figure 2B), locked to the stimulus onset (left panels) and locked to the response time (right panels).

**Figure 2.**
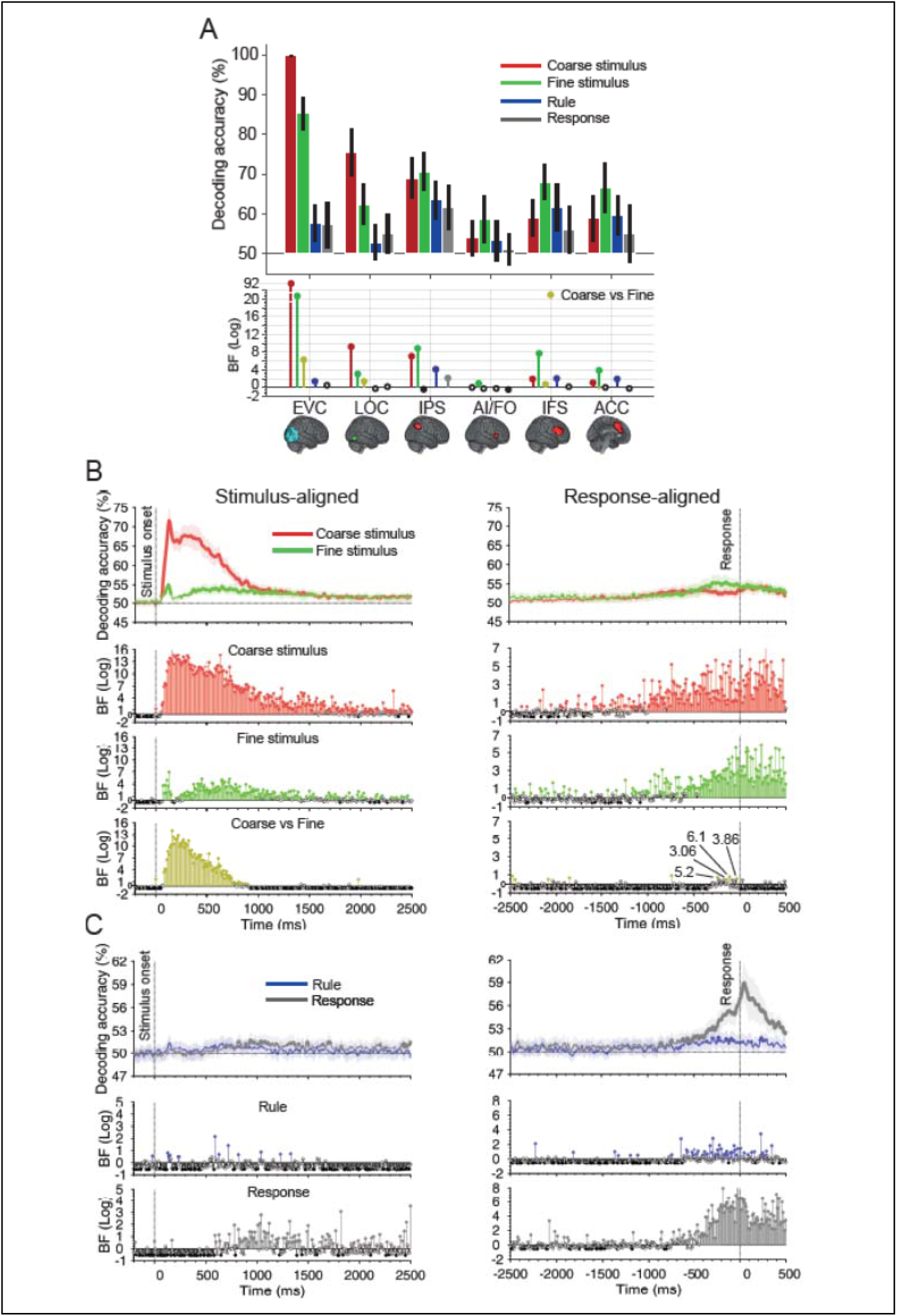
Decoding of different types of information from fMRI ROIs and MEG time points. (A) Average fMRI decoding (error bars = 95% confidence interval across participants) for every ROI. The theoretical chance-level decoding is 50%. Bottom panel: Bayesian evidence for above-chance decoding for each condition (relative to chance) and difference between coarse and fine stimulus information decoding (yellow), BF: Filled coloured circles show evidence (BF>3) for the alternative hypothesis, and filled black circles reflect evidence for the null hypothesis (BF<1/3). Empty circles indicate insufficient evidence (1/3<BF<3). BF were calculated using Bayesian t-tests. (B) Stimulus- and response-aligned decoding of stimulus information across time in MEG data. Thick lines show the average across participants (shading 95% confidence intervals), double-thickened lines show time points with evidence (BF>3) for above-chance decoding. BFs are shown every 10 ms (rather than 5 ms) for clearer illustration. Horizontal dotted line refers to theoretical chance-level decoding (50%). Vertical dotted lines indicate critical times in the trial. Bottom panels should be read as in (A). (C) Same as (B) for rule and response information. Fine stimulus information, rule and responses were reproduced from Robinson et al., 2022. Note the different scales of the BF panels.

We quantified the strength of representations about four orthogonal aspects of information using decoding (see *Methods,* and Figure 1A), namely coarse and fine stimulus information, rule information and response information. Specifically, for stimulus information, we looked at two possible levels of required precision in representations of stimulus location. To quantify *coarse* stimulus information, we decoded information about side of the visual hemifield where the stimulus was presented. To quantify *fine* stimulus information, we decoded information about the precise location of the stimulus within each visual hemifield. We decoded the coarse and fine stimulus information and compared them, to test whether these different aspects of the visual representation are encoded and transferred in a similar or different manner across the brain, testing the role of MDN in prioritising perceptually difficult vs. easy aspects of stimulus information (Woolgar et al., 2011; Woolgar et al., 2015b). For rule information, we decoded the information about the rule (Rule 1 or 2). For response information, we decoded trials where inner vs. outer fingers were pressed.

#### ROI-based fMRI decoding

In EVC, we observed strong representation of stimulus information (Figure 2A). The classifier was able to distinguish both the coarse (red bars) and fine (green bars) stimulus information (coarse stimulus information BFs for EVC=1.4e92, LOC=1.6e9 and fine stimulus information BFs for EVC=4.6e20, LOC=1.1e3), and there was evidence that coding of coarse stimulus information was stronger than fine stimulus information (BF=1.7e6), reflecting the magnitude of the visual differences in coarse and fine stimulus comparisons. There was also evidence that the EVC represented rule information (BF=23) but there was insufficient evidence to determine whether rule was presented in the LOC (BF=0.48), or whether response information was represented in either ROI (BFs for EVC=3, LOC=1.4).

The MDN, on the other hand, exhibited a different pattern of information coding. There was evidence for coding of both coarse and fine stimulus information in all ROIs (coarse stimulus information BFs for IPS=1.1e7, IFS =75, ACC=12, and fine stimulus information BFs for IPS=6.6e8, AI/FO=7.8, IFS =46e6, ACC=6.6e3), except for the AI/FO (insufficient evidence for coarse; BF=0.92). However, the pattern for stimulus information was opposite to the EVC, with coding of fine stimulus information being numerically stronger than coding of coarse information in all regions, and statistical evidence for a difference in the IFS (BF=5). There was evidence that rules were represented across all the MDN (BFs for IPS=1.2e4, IFS =99, ACC=74) except for the AI/FO (insufficient evidence; BF=0.55). Only the IPS held information about response (BFs for IPS=130, AI/FO=0.31, IFS =1.4, ACC=0.55). The CO sub-network (AI/FO and ACC) numerically showed lower levels of information compared to the FP sub-network (IFS and IPS) of the MDN.

#### Time-resolved MEG decoding

In MEG, we used data from a previous study using the same paradigm in an independent group of participants (Robinson et al., 2022). As in that study, we used the stimulus-locked and response-locked analyses to examine different aspects of information processing: whereas stimulus-driven signals are likely to be strongest when locked to stimulus onset, rule application and decision making requires computation based on the cue and can take variable amounts of time on different trials, and so is likely to be more evident when we align the neural signals based on the time of response. Here we examine coding of a new aspect of the task (coarse stimulus information). We also re-ran the time-resolved decoding of fine stimulus, rule and response coding reported in (Robinson et al., 2022). We reproduce the results here for ease of comparison to the coarse stimulus information and the fMRI decoding results.

Coarse and fine stimulus information were present (BFs>3) when the signals were aligned to stimulus onset from 55ms and 65ms post-stimulus onset, respectively, both peaking at 130 ms (Figure 2B, left). Both types of visual information were sustained until after 1000 ms with evidence (BFs>3) for stronger coarse than fine stimulus information until 800 ms post stimulus onset. In contrast, when the signals were aligned to response, there was evidence (BFs>3) for greater coding of *fine* than coarse stimulus information between 300 to 100 ms before the response (Figure 2B, right).

For rule coding, we reproduce here the results reported in (Robinson et al., 2022). The *stimulus-aligned* analysis showed only a few sparse time points with evidence (BFs>3) for above-chance rule information after the stimulus onset (Figure 2C, left, blue), but *response-aligned* analyses showed numerous consecutive time points with evidence (BFs>3) before and around the time of response (from -500 to 500 ms post response time; Figure 2C, right, blue).

We also reproduce here the results for response coding reported in (Robinson et al., 2022). There was little response information in the *stimulus-aligned* analysis (Figure 2C, left, grey) but when the data were *response-aligned*, there was evidence (BFs>3) for response information around the time response starting from around -700ms and lasting until the end of the analysis window (Figure 2C, right, grey).

Comparing the fMRI and MEG data qualitatively, we observe that the pattern of information coding in the EVC, with stronger coding of coarse relative to fine stimulus information, and relatively weak evidence for coding of rules and response, mirrors the data from earlier timepoints in the MEG. Conversely, data from several MD regions, reflecting stronger decoding of fine compared to coarse stimulus information, and task rules, more closely mirrors the MEG data from later timepoints. Next, we sought to formalise these observations and study the dynamics of information coding in each region using model-based MEG-fMRI fusion.

### Analysis 2: spatiotemporal dynamics of information encoding (fMRI-MEG fusion)

We used model-based fusion of fMRI and MEG representations (Hebart et al., 2017, Moerel et al., 2024) to quantify when, and in which ROI, the structure of representational space in the two modalities correlated with each other and with a theoretical model that captured the coarse stimulus, fine stimulus, rule and response information (Figure 3A). This allowed us to quantify, for each region, the timecourse with which different aspects of the task were represented. The commonality profiles in general followed the MEG decoding profiles with stimulus information more aligned to the stimulus onset (Figure 3B, left) and the rule and response commonalities peaking before and around the response (Figure 3B, right). Significant (p<0.05; random permutation testing corrected for multiple comparisons) commonalities are shown by thickened lines. We used a permutation approach rather than Bayes factor analysis here as there was no way to *a priori* set the chance level of commonality.

**Figure 3.**
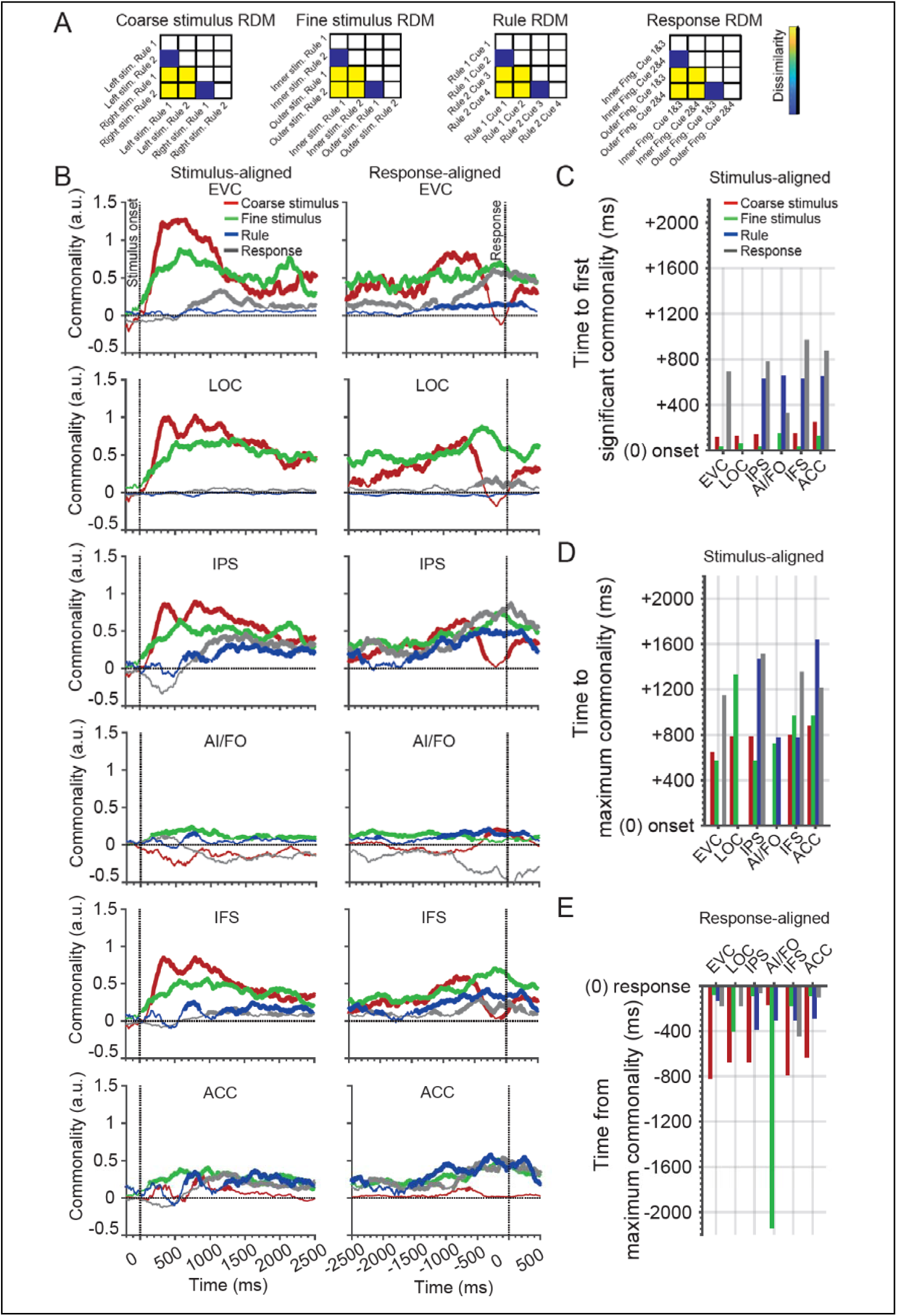
Commonalities between fMRI, MEG and models. (A) Four distinct neural Representational Dissimilarity Matrices (RDMs) were constructed for each fMRI ROI and each MEG time point by different arrangements of stimuli, rules, cues and responses, which allowed us to evaluate the commonality between fMRI and MEG representations and target models (here we are showing the models rather than neural RDMs). Note that the rows and columns of these matrices are different for each task feature (see *methods*). Model RDMs had identical arrangements of conditions to the neural RDMs, but only included binary values (1 for high and 0.5 for low dissimilarity). (B) Commonality time courses aligned to the stimulus-onset (left column) and response (right column) times. Thickened lines indicate significant commonalities (p<0.05; one-tailed random permutation testing; corrected for multiple comparisons across time). Note that only positive commonalities are interpretable as they reflect the correlation between modalities. Horizontal dotted line indicates no commonality and negative commonalities are not meaningful. Vertical dotted lines indicate critical times in the trial. (C-E) Quantitative commonality metrics: Time from the stimulus onset to the first significant (C) and to maximum (D) commonalities, and the time from the maximum commonality to the response (E), extracted from the commonality time courses for each information type. Bars are not shown where commonality traces did not reach significance.

In the *stimulus-aligned* analysis, there was numerically higher commonality for coarse than fine stimulus information in EVC, LOC, IPS and IFS. However, in the *response-aligned* analysis the commonality was numerically higher for fine than coarse stimulus information in all these regions around the time of response. Thus, the general pattern seen in the MEG data above was born out in this analysis again and could be ascribed to the responses of the EVC and MDN. However, the CO sub-network (AI/FO and ACC) showed a different pattern to the FP sub-network of the MDN. In the CO sub-network, there was little commonality overall, but what coding there was showed a more sustained pattern than the FP sub-network. In particular, as opposed to the FP sub-network, the AI/FO showed significant coding of only the *fine* stimulus information around the time of stimulus representation, and coding of only the coarse stimulus information around the time of the response, at around the time that the FP sub-network stopped coding the coarse information. The ACC showed a sustained representation of only the fine visual information both at after stimulus onset (*stimulus-aligned*) and around the time of the response (*response-aligned*). This may suggest a different role for the AI/FO and ACC relative to the more lateral and dorsal FP sub-network of the MDN in stimulus processing, as supported by previous studies (Dosenbach et al., 2006; Crittenden et al., 2016).

In the stimulus-aligned analysis, rule information first appeared later in time than the visual information (after ∼700 ms) and was seen in all MDN but no visual areas (Figure 3B, left). In the *response-aligned* analysis, all nodes of the MDN showed the rule information from ∼-2500 ms before the response (ramping up and peaking at ∼-500 ms before the response), the EVC showed rule information only appearing later, ∼-1000 ms before the response (Figure 3B, right). Therefore, the significant rule coding in the EVC in the fMRI data may have arisen at later timepoints, perhaps reflecting feedback.

Not surprisingly, the rising temporal profile of response information was more clearly observed in *response-aligned* analysis, which peaked around the time of response in all areas except AI/FO, which did not show response coding.

The commonality results suggest distinct temporal profiles of information encoding between the different types of information and brain regions. To quantify the comparison between different types of information, we calculated “time to first significant commonality”, “time to maximum commonality” and “time from maximum commonality” for each region and information aspect (see *Methods*; Figure 3C-E). These are only single data values as fusion was performed using group-averaged data across modalities. In *stimulus-aligned* data, the time to the first significant commonality was numerically slightly shorter for the fine (green bars) than coarse (red bars) stimulus information across all areas (except for AI/FO where coarse stimulus information was never significant so we do not have a data point; Figure 3C). However, the times to maximum commonality for the two types of stimulus information were similar (for all regions except LOC where coarse information was first; Figure 3D). In *response-aligned* data, commonalities also reached their peaks earlier for coarse relative to fine stimulus information, across all areas except AI/FO, which, as noted above showed a low but sustained representation of the fine stimulus throughout the analysis window (Figure 3E).

Comparing between regions, relative to the stimulus onset, the time to the *maximum* commonality increased from posterior visual areas to anterior nodes of the MDN for the coarse (EVC = 645 ms; LOC = 785 ms; IPS = 785 ms; AI/FO = N/A; IFS = 795 ms; ACC = 880 ms) and fine (EVC = 570 ms; IPS = 570 ms; AI/FO = 725; IFS = 970 ms; ACC = 970 ms; LOC = 1330 ms) stimulus information. This suggests potential feed-forward flow of stimulus information from EVC to MDN (not including LOC, for which the time course suggests it might have been by-passed).

In *response-aligned* data, the time from maximum commonality (to response) for rule information suggests that, relative to response times, rule information was dominantly encoded by the posterior MDN areas followed by more anterior areas, and finally appeared in the EVC just prior to the response being actually made (LOC = N/A; IPS = -385 ms; AI/FO = -305; IFS = -305 ms; ACC = -285 ms; EVC = -130 ms).

The time from maximum commonality to response for response information did not show a clear pattern but was shortest in the IPS area (AI/FO = 1425 ms; IFS = 445 ms; EVC = 175 ms; LOC =175; ACC = 100 ms; IPS = 60 ms) suggesting that this area might be one of the critical ROIs in the MDN to support motor responses.

### Analysis 3: evaluating flow of information across the brain using Fusion-RCA

Finally, we tested for statistical relationships between the dynamic representational profiles of the different regions using an adaptation of Granger causality (Barnett and Seth, 2014), to quantify the potential exchange of information between them and characterise how information was circulated within and beyond the MDN. To do so, we applied Granger-causality analysis to the commonality indices from different ROIs to evaluate their statistical dependency in time, a new method that we call *Fusion-RCA* (Figure 4A). This analysis goes beyond co-variation of activations or representational structures (Goddard et al., 2016; Anzellotti and Coutanche, 2018; Karimi-Rouzbahani et al., 2022) and considers the relationships between the commonality time courses obtained for each ROI in the fusion analysis. We evaluated the flow of information separately for incoming information to and outgoing information from any of the 6 target ROI using multivariate (multi-ROI) Granger causality (Barnett and Seth, 2014; see *Methods*).

**Figure 4.**
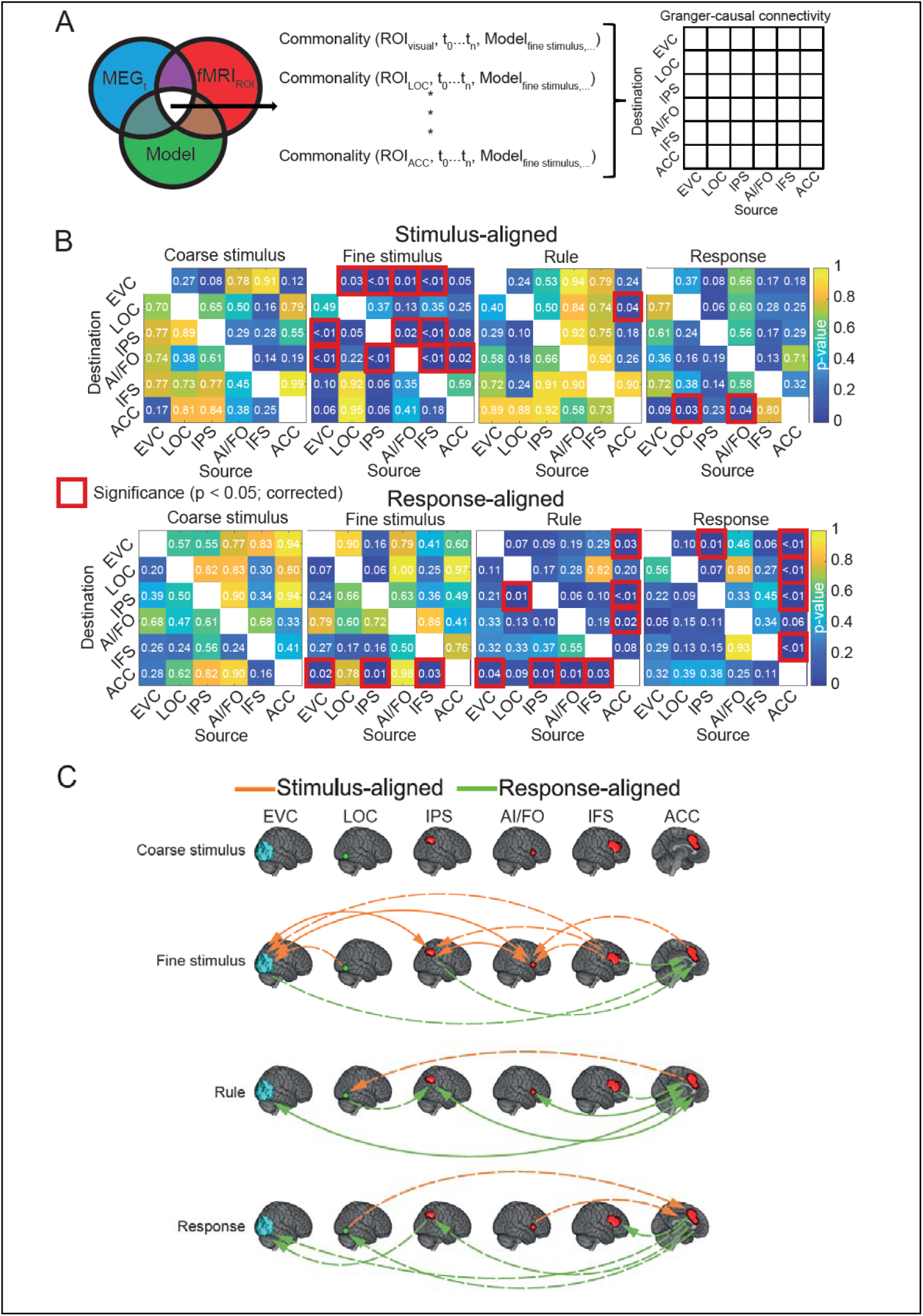
Fusion-based analysis of connectivity. (A) RDMs from every fMRI Region of Interest (ROI), every MEG time point, and every model were entered into the commonality index equation (equation (1), see *Methods*), and then the resultant commonality time series were fed to the multivariate Granger causality (MVGC) toolbox to determine potential flow of information between all six ROIs in both directions. (B) *p*-values of connectivity between every pair of ROIs in *stimulus* (upper) and *response* (lower)-*aligned* analyses. *p*-values were obtained using asymptotic null distribution corrected for multiple comparisons for the number of time samples and ROIs. Significant p-values (indicative of information exchange; <0.05) are shown by red boxes. (C) Information transfer diagrams, visualising the results in (B). Arrows show the existence and the direction of information flow across ROIs in the *stimulus*- (orange) and *response* (green)-*aligned* analysis. Unidirectional connections are shown using dashed lines and bidirectional connections are shown using solid lines.

The results of the Fusion-RCA are shown in Figure 4B. For interpretation, we focus on the significant results diagrammatically shown in Figure 4C. For *coarse* stimulus information, we found no significant (p<0.05; corrected for multiple comparisons; Figure 4C) flows across any of the ROIs, regardless of either alignment to the stimulus onset or the response time. This suggests that while coarse stimulus information was strongly represented in visual areas, it was not significantly exchanged with and within the MDN.

In contrast, for *fine* stimulus information (Figure 4C), we observed abundant flow between all ROIs, suggesting considerable exchange and circulation of information pertaining to this more challenging aspect of visual information. The *stimulus-aligned* analysis suggested a critical role for the EVC as a receiver of fine-grained visual information and AI/FO as the most active node of the MDN. We observed information coming from LOC, IFS, IPS and AI/FO to the EVC and output from this region to the IPS and AI/FO. AI/FO was the only MDN region exchanging information with all other MDN regions. In the *response-aligned* analysis, the ACC seemed to be the apex of processing, receiving fine stimulus information from EVC, IPS and IFS areas. In summary, all MDN areas took part in circulating the fine stimulus information. Upon stimulus onset, information from EVC flowed into the MDN via the IPS and AI/FO nodes and was circulated back to the EVC from all lateral MDN regions. Before the response, information was fed to the medial ACC region.

As expected, flow of *rule* information was most visible in the *response-aligned* data. Here we again observed a critical role of the ACC in receiving rule information (from EVC, and all MDN regions) and feeding rule information elsewhere in the system (in EVC, IPS, and AI/FO regions), while those other MDN regions did not appear to exchange rule information directly.

In the exchange of *response* information, ACC again played a central role, receiving input (stimulus-aligned) from LOC and AI/FO and driving response information (*response-aligned*) into almost all other regions. There was also exchange of response information from IPS to EVC, potentially accounting for the representation of response information in these regions at late timepoints (c.f., Figure 3B, right).

Together, these results showed that not all types of information were exchanged as strongly across the MDN. Whereas there was no significant flow of (perceptually easy) coarse stimulus information, there was significant circulation of (perceptually hard) fine stimulus information, rule and response information between nodes of the MDN and visual areas. These support a role for the MDN in connecting distinct compartments of the brain depending on task demands (Duncan et al., 2020).

Moreover, we saw that sub-regions of the MDN play distinct roles in transferring different types of information within the same trial. The fusion-based connectivity analysis allowed us to characterise the direction and the content of the transferred information simultaneously which might be misinterpreted if we look at the fusion commonality traces alone. For example, while the temporal dynamics of the commonality traces appeared to emphasise potential feedback of rule information from the MDN to EVC, the connectivity analyses clarified that rule exchange with EVC was bidirectional and mainly involved the ACC MDN region. Finally, we saw a critical role for the ACC in exchanging different types of information including fine stimulus, rule and response information. We also observed that AI/FO also exchanged information with ACC for all these aspects of information suggesting their close functional cooperation and potential distinction from the FP sub-network of the MDN (Crittenden et al., 2016).

## Discussion

The multiple demand network has been suggested to be a potential candidate for connecting distant and functionally distinct areas of the brain to construct “integrated intelligence from distributed brain activity” (Duncan et al., 2020, p.838). However, the temporal dynamics of information coding and flow in the MDN and the role of distinct nodes of the MDN in information transfer have been challenging to investigate, especially in humans. Here we used MVPA in spatially (fMRI) and temporally (MEG) high-resolution neuroimaging modalities and showed that different parts of the MDN represent distinct types of information to varying degrees. We found that coding in the MDN emphasised the more fine-grained aspects of visual stimuli, despite this being a smaller physical visual signal and one that was coded less strongly than the large coarse grain stimulus distinctions in visual regions. Then we used the MEG data to show distinct temporal profiles for coarse- and fine-grained stimulus information. Together these results suggested time-varying involvement of MDN and other brain areas in information coding. To formalise this, we fused fMRI and MEG data and showed that coarse- and fine-grained stimulus information had distinct time courses of coding in all areas of the MDN. This provided evidence for the highly dynamic nature of information coding in the MDN depending on stimulus type and difficulty. Finally, we developed a novel fusion-based connectivity analysis which allowed us to trace information exchange within and beyond the MD system. This revealed network-wide engagement and circulation of the challenging aspects of stimulus processing (fine stimulus information), in the absence of information exchange about the large physical (coarse) differences. It also revealed the critical role of the EVC and the ACC in the analysis of fine stimulus information, with the ACC also critical for receiving and distributing rule and response information. This provides unprecedented resolution on the distinct roles of different MDN regions in the encoding and exchange of multiple types of task information.

Human fMRI studies have shown for two decades that the nodes of the MDN become active and encode task-relevant (and usually challenging) aspects of information across a variety of tasks (Duncan and Owen, 2000; Duncan, 2010). Due to the temporal resolution of fMRI, however, researchers have been unable to determine whether this apparently simultaneous activation pattern was due to the nodes of the MDN having similar temporal profiles (and therefore doing similar things), or a lack of temporal resolution. Here, we combined fMRI with MEG and found detectable differences between the temporal dynamics of information encoding across the nodes of the MDN for distinct aspects of information. This detailed picture of the dynamics of information processing across the MDN supported the possibility of the MDN’s role in brain-wide information exchange through information integration and propagation. Our findings are consistent with an fMRI study that provided evidence for the feeding of sensory-related activations into adjacent nodes of the MDN (Assem et al., 2021). However, their univariate activity-based analysis did not evaluate the information content potentially fed into the MDN, but together our results are in line with the hypothesis about the role of the MDN. Similarly, a recent study also found stimulus category information in parietal and frontal brain areas near the core MDN suggesting potential flow through the MDN, although they did not quantitatively evaluate connectivity (Shashidhara et al., 2024).

We observed that not all regions of the MDN encoded and transferred information similarly. Specifically, we observed numerically lower (c.f., Figure 2A) and more sustained (c.f., Figure 3B) information coding in the cingulo-opercular (i.e., AI/FO and ACC) than lateral frontoparietal (i.e., IPS and IFS) sub-networks of the MDN. We also observed that ACC played a central role in transferring information across the MDN and beyond while consistently exchanging information with AI/FO about different aspects of the task (c.f., Figure 4C). These observations align with the distinctions made between the CO and FP sub-networks within the MDN, where studies have reported not only better encoding of task-relevant information in CO than FP but also more pronounced within- than cross-sub-network functional connectivity (Crittenden et al., 2016). The sustained and less variable encoding of information in the CO than FP sub-network observed here (c.f., Figure 3B) aligns with previous studies which split the MDN into CO and FP sub-networks (Dosenbach et al., 2006) based on findings that FP activations changed on a trial-by-trial basis whereas the FP sub-network showed sustained activations over blocks of trials (Visscher et al., 2003). Among the MDN regions, we observed a central role for the ACC in information exchange with other brain areas. This may reflect its critical role in different aspects of human cognition and executive control (Carter et al., 1999). Specifically, the dominantly cognitive ACC region targeted in the present study has been shown to be involved in attention and motor control and especially activated in cognitively demanding tasks where stimulus and response selections were difficult such as in Stroop tasks (see Bush et al., 2000 for a review).

We have previously shown a direct relationship between (stimulus and rule) difficulty and the level of information coding in the MDN (Woolgar et al., 2011a; Woolgar et al., 2015a; Woolgar et al., 2015b). The current results extend previous observations and highlight the role of the MDN in the encoding of the difficult aspects of stimuli without explicit cueing in the experimental design. Specifically, our previous studies compared distinct sets of stimuli (e.g., with vs. without noise (Woolgar et al., 2015b), or at different eccentricities (Woolgar et al., 2011a)) in the hard vs. easy conditions to show the preference of the MDN in coding of hard vs. easy stimulus conditions. As the two stimulus sets differed (in terms of noise or eccentricities), their comparison might have been affected by stimulus-related differences rather than difficulty only. The tasks also had a blocked design with each block containing either the easy or hard condition. Therefore, it might have been the case that the higher encoding in the MDN in hard vs. easy blocks reflected a general effort signal initiated by the cue at the beginning of each block to modulate the representations of information. In the current study, on the other hand, we used four stimuli in an event-related design and quantified the level of hard vs. easy aspect of the same stimuli in an orthogonal decoding fashion. Specifically, all four stimuli were included in the decoding whether decoding the coarse (left vs. right side) or fine (inner vs. outer) aspect of the stimulus. Critically, participants needed to process both aspects of the stimulus information on every trial for every single stimulus to successfully perform the task, but probably had more difficulty telling if a stimulus was inner vs. outer in the visual field (fine information) than if on the left vs. right side (coarse information). Therefore, the easy vs. hard aspect of stimulus discrimination existed implicitly on every trial rather than being cued. Our results showed that the MDN seem to automatically (without any explicit cues about the difficulty of the information and the effort exerted by the participant) provide an enhanced representation for the more fine-grained aspect of the visual input which was not as well represented in the visual system. This is consistent with the role of the MDN in processing difficult aspects of the task.

One novelty of the current work is the development of a fusion-based connectivity analysis, *Fusion-RCA*. This connectivity analysis provides several advantages over conventional connectivity analyses. First, it follows the recent shift from univariate to multivariate connectivity analyses (Coutanche and Thompson-Schill, 2013; Geerligs and Henson, 2016; Goddard et al., 2016; Anzellotti and Coutanche, 2018; Basti et al., 2020; Karimi-Rouzbahani et al., 2021a; Karimi-Rouzbahani et al., 2022). Specifically, conventional univariate connectivity analyses, which evaluate statistical dependency across individual sensors, are prone to missing statistical relationships/connectivity reflected in low-amplitude activity (Anzellotti and Coutanche, 2018), as well as high-dimensional patterns across multiple sensors (Geerligs and Henson, 2016; Basti et al., 2020). On the other hand, multivariate (Rahimi et al., 2022), also called multi-dimensional (Basti et al., 2020) and representational (Karimi-Rouzbahani et al., 2022) connectivity methods, are less affected by absolute activity values and can detect high-dimensional (across many sensors/voxels/electrodes) statistical relationships. Second, while conventional univariate analyses evaluate connectivity by quantifying the statistical relationship between activations, Fusion-RCA quantifies information transfer and provides insights about the content of the transferred information. Third, it provides a higher spatial resolution than conventional (source-space) MEG (>1.0 cm; Hauk et al., 2011) as it uses the voxel-level (here 3.0 mm^3^) fMRI representations. It also provides a higher temporal resolution than comparable connectivity analyses in fMRI such as informational connectivity (Coutanche and Thompson-Schill, 2013) or jack-knife-based representational connectivity (Coutanche et al., 2020) as it utilises the millisecond scale (here 5ms) time-resolved representations from MEG. This additional spatiotemporal resolution can be crucial for characterising highly dynamical systems such as the MDN whose nodes had shown very similar activations in previous fMRI studies (Fedorenko et al., 2013; Duncan et al., 2020). Without fusion analysis, the temporal dynamics of information encoding across multiple brain locations would require high spatiotemporal resolution data from multiple regions in animal models (Panichello and Buschman, 2021) or patients implanted for intracranial recordings (e.g., Karimi-Rouzbahani and McGonigal, 2024). Importantly, this fusion-based connectivity analysis is not limited to the current study but can be used to approach many questions in cognitive neuroscience where high spatiotemporal resolution is necessary.

An important point about representation- and/or information-based *connectivity* is its distinction from information *representation/encoding* within individual ROIs. Specifically, a given pair of ROIs might encode similar information (above chance) but not interact with each other or they might exchange a specific aspect of information but do not encode it significantly (Anzellotti and Coutanche, 2018). Therefore, inference about potential connectivity should not be based on decoding profiles or fusion commonalities *per se* (Cichy et al., 2014; Mohsenzadeh et al., 2018). For example, we observed that although rule information was initially maximally encoded across the MDN followed by the EVC, connectivity analysis suggested bidirectional feed-forward and feedback flows of rule information around the time of response. While the feed-forward flow of rule information may be due to rule information needing to be extracted from the cues which are provided visually, the feedback of rule may also enhance the relevant visual information in sensory areas. So, this extraction process necessitates involvement of visual followed by higher-order cognitive areas. Fusion-RCA proposes that connectivity analysis can be built on the decoding profiles (in MEG or fMRI separately) or commonality indices which have been increasingly used in the field to provide a more detailed picture of the potential interaction of information between brain areas (Cichy et al., 2014; Mohsenzadeh et al., 2018; Hebart et al., 2017; Moerel et al., 2024).

While there are benefits in using fMRI-MEG fusion and Fusion-RCA for obtaining insights about the spatiotemporal dynamics of information encoding and connectivity, there are also limitations (Cichy et al., 2020). First, fusion is only useful when effects are present in both modalities; no additional effect will be generated through fusion. If effects appear in one imaging modality but not the other, they will be diminished or not detected. Second, as weak effects might be further weakened in fusion through imperfect correlation across modalities, fusion is more suited for medium to large effects that are clear in both modalities. This is particularly the case here, where the fMRI and MEG data come from separate groups of participants. Third, the signal-to-noise ratio of the information can also be an important factor when interpreting peaks of commonalities in fusion. For example, it is assumed that frontal MD areas receive visual signals later than visual areas. However, a large early peak of visual signals might cause a larger commonality index in that same MD area followed by a smaller peak which is caused by when visual signals reach the MD area. Therefore, peaks of commonality might sometimes be hard to interpret for extracting information transfer. Finally, while we did not delve into potential *transformation* of information across the brain in this work, all the significant connections involved some levels of information transformation across areas. In other words, if we had two fMRI ROIs with identical representations, our Fusion-RCA would not detect any difference in their commonality time courses and no causal relationship would have been found, because the correlation of two ROIs with identical representations with one MEG data would result in identical correlations – no inter-area difference. Therefore, it is important to note that all the detected transferred information have also been transformed from the source to destination. Therefore, while fusion and fusion-based analyses can provide additional insights into potential exchange of information in the brain, it is crucial to keep in mind these considerations when interpreting their results.

Because of previous challenges with temporal and spatial resolution in neural recording modalities, there have not yet been strong predictions about the specific involvement of different MDN nodes in the flow of information in an ordered manner. However, here we demonstrated that approaching the problem in a data-driven way allows for new insights. For example, we identified the apparently critical role of the ACC in information transfer across the MDN. Fusion-RCA method can also be used for future studies where the content of the transferred information is unknown. In the case of the MDN, for example, it would also be interesting to investigate how the patterns of information exchange change under different circumstances, such as during memory recall, or when directed selective attention is applied.

In conclusion, this study fused fMRI and MEG, and developed a novel fusion-based connectivity analysis, to provide new insights into the temporal dynamics of information encoding and transfer across and beyond the MDN. We showed that the MDN’s involvement varies across different aspects of the task including higher involvement and encoding of difficult vs. easy aspect of stimulus information. We also found subtle but detectable temporal differences in the time course of information encoding across the MDN for distinct information types, suggestive of a distinction between the frontoparietal (FP) and cingulo-opercular (CO) sub-networks of the MDN which played different roles in information exchange between the MDN and visual areas during the task. We found a central role for the ACC in information transfer in our experiment, presumably reflecting its multifunctional role in supporting different aspects of human cognition and behaviour. This work provides new insights about the potential role of the MDN as one of the main candidate mechanisms supporting complex task performance, and introduces Fusion-RCA, a novel connectivity method which can be used to address many questions about cognition.

## Methods

We collected fMRI data using the same paradigm for which we had previously collected MEG data (Robinson et al., 2022), with new participants. The paradigm is designed to investigate the encoding of stimulus, rule and response in a stimulus-response mapping task and has shown a strong link between behavioural performance and neural coding of information (Woolgar et al., 2019). Specifically, at around the time of response, the coding of stimulus representations in the MDN predicted whether participants would give a correct or incorrect response (Robinson et al., 2022). We combined this MEG dataset with our new fMRI dataset to explore the role of the MDN in encoding and integrating task information for flexible behaviour.

### Participants

30 (14 female, 16 male; age: mean = 24.4 years, sd = 2.1) participants were recruited from Macquarie University volunteer panel for the fMRI study. No fMRI participant had participated in the previous MEG study (see Robinson et al. 2022, for full methodological details of the MEG study). Participants were right-handed and had normal or corrected to normal (through contact lenses) vision. Participants gave written informed consent for both a behavioural training of 1 hour prior to scanning (reimbursed AU$15) and the 2 hour fMRI experiment (including setup; reimbursed AU$40). The study was approved by the ethics committee of Macquarie University.

### Task design

We collected fMRI data while participants performed a visual stimulus-response mapping task (Robinson et al., 2022) designed to tease apart stimulus, cue, rule and response encoding in the brain. The task involved pressing of one of four buttons according to the position of a visual stimulus and one of two memorised stimulus-response mapping rules (Figure 1A). Each trial started with the presentation of a grey fixation square for 500 ms followed by the stimulus for another 500 ms after which the participants could provide a response (Figure 1B). The response screen lasted for 4000 ms or until the response was made, whichever happened first.

In the initial behavioural training session, participants had to learn two set of mappings between stimuli and the buttons (rules), as indicated by the colour cue on the centre of the screen. Each rule was indicated by two cue colours (giving a total of 4 possible cue colours: two indicated that Rule 1 was to be applied, and the other two indicated that Rule 2 was to be applied) to allow us to see the effect of cue colour on rule processing, and to avoid the classifiers using cue colour to distinguish the categories. Specific cue colour-rule associations were randomised and counterbalanced across participants. Participants responded using the four fingers of their right hand.

Stimuli were presented and responses collected using MATLAB Psychtoolbox (Brainard, 1997). Our training session was identical to the one used in the MEG study (Robinson et al., 2022). Briefly, the task started from easier versions, in which stimuli were presented in non-overlapping positions in earlier blocks, to the final version, with overlapping stimuli, like the version used in the MRI scanner. Rules were trained one by one in the first two blocks, each block containing 32 trials. Participants needed to reach 60% accuracy to continue to the subsequent block, otherwise the block would repeat until they reached the threshold or they were excluded and replaced if they did not reach it after 5 consecutive blocks. Participants did an average of 8.61 (sd = 2.46) training blocks to finish the training session successfully.

### Apparatus

FMRI scans were acquired using a Siemens 3 T Verio scanner with 32-channel head coil, at the Macquarie Medical Imaging centre, Macquarie University Hospital, Sydney, Australia. We used a high resolution interleaved ascending T2*-weighted echo planar imaging (EPI) acquisition sequence with the following parameters: repetition time (TR), 2000 ms; echo time (TE), 30 ms; 36 slices of 3.0 mm slice thickness with a 0.366 mm interslice gap; in-plane resolution, 3.0 × 3.0 mm; field of view, 126 mm. We also acquired T1-weighted MPRAGE structural images for all participants (non-selective inversion recovery, resolution 1.0 × 1.0 × 1.0 mm). The session began with the collection of a T1 image which lasted for about 5 minutes followed by EPI acquisition for the blocks of experimental trials.

Each block started with the presentation of a rule screen which showed the four cue conditions, 2 per rule, for 10 seconds, followed by the experimental trials. We acquired 6 scanning runs, each consisting of two blocks. Each run lasted for ∼ 8 minutes after which the data were saved and scanning restarted. Within each block, there were 80 trials presented in pseudorandom order such that all stimulus configurations (4 cue colours * 4 stimuli) appeared five times with equal probability within each block. Stimuli were presented on an LCD screen at the back of the scanner, and participants could see the LCD through a head-coil mounted mirror.

### Pre-processing

We pre-processed the fMRI data using SPM 12 (Wellcome Department of Imaging Neuroscience, www.fil.ion.ucl.ac.uk/spm) in Matlab 2018a. DICOM data were converted to NIFTII format. Functional images were slice-time corrected and realigned to the first functional scan in the run. They were then smoothed slightly (4 mm FWHM Gaussian kernel) to increase the signal-to-noise ratio as in previous work (Woolgar et al., 2015b; Jackson et al., 2017). Structural images were co-registered to the mean functional images and normalised to the MNI152 space (McConnell Brain Imaging Centre, Montreal Neurological Institute) to derive normalisation parameters that we later used to define Regions of Interest for individual participants.

The MEG pre-processing procedure is explained in the MEG study in detail (Robinson et al., 2022). Briefly, the signals were band-passed (0.03-200Hz), notch-filtered (50Hz) and down-sampled to 200 Hz with no additional pre-processing steps. Data were cut into *stimulus-aligned* epochs from -200 before to 2500 ms after stimulus onset and *response-aligned* epochs from -2500 to 500 ms after response. Signals are analysed in the 160-channel sensor space.

### Regions of Interest (ROIs)

Template space MDN ROIs were taken from our previous work (Woolgar et al., 2015b) where they were defined based on their activity in a wide range of cognitive tasks (Duncan, 2010). These consist of left and right inferior frontal sulcus (IFS; centre of mass +/−38 26 24, volume 17000 mm^3^); left and right anterior insula/frontal operculum (AI/FO; +/−35 19 3, 3000 mm^3^); left and right intraparietal sulcus (IPS; +/−35 −58 41, 7000 mm^3^) and the dorsal anterior cingulate area (ACC; +/−0 23 39, 21000 mm^3^).

The visual areas were obtained from the Brodmann template provided with MRIcro (Rorden and Brett, 2000): Brodmann area 17/18 (EVC; −13 −81 3, 16 −79 3, 54000 mm^3^). The lateral occipital complex (LOC; +/− 40 − 70 – 9, 41800 mm^3^) was defined from a prior review of 8 imaging studies reporting greater activation for objects compared to scrambled shapes or textures (Thompson and Duncan, 2009). ROIs were defined based on a subjects-averaged localiser data from a previous study where we contrasted the areas which responded more strongly to pictures of natural/man-made objects than to scrambled versions of the same objects (Jackson et al., 2017). Our LOC ROIs fell very close to anatomical LOC coordinates from previous studies (Grill-Spector et al., 2000, Grill-Spector et al., 1999). All ROIs were defined and analysed in each hemisphere separately, but decoding results were averaged over hemispheres for inference.

### First-level model

To estimate the activation patterns (beta values) for each condition and each area we used a General Linear Model (GLM), convolving event regressors with the duration of the trial reaction time (Grinband et al., 2008) with the first-order hemodynamic response of SPM. For MVPA, we used a total of 8 regressors representing *coarse* stimulus information (the two stimuli on the left vs. the two on the right side of space), *fine* stimulus information (the two stimuli located in the inner vs. outer peripheral positions relative to the central fixation square), *rule* (rule 1 vs. rule 2) and *response* (inner vs. outer buttons). Each trial contributed to the estimation of 4 of these regressors. For RSA analysis, we ran 4 separate sets of GLMs on the data to maximise the power for the desired information to avoid subtle information (e.g., rule) being dominated by stronger information (e.g., coarse stimulus)^1^ and to have enough regressors for RSA (see Figure 3A). Specifically, to construct a 4×4 representational dissimilarity matrix (RDM) for coarse stimulus information, we included two regressors for the coarse stimulus information (left vs. right) and two regressors for the rule information (rule 1 vs. rule 2). To construct the 4×4 RDM for fine stimulus information, we included two regressors for the fine stimulus information (inner vs. outer) and two regressors for the rule information (rule 1 vs. rule 2). To construct the 4×4 RDM for rule information, we included four regressors for the rule information (all four cue colours). Finally, to construct the 4×4 RDM for response information, we included two regressors for the response information (inner vs. outer buttons) and two regressors for the cue colours information collapsing across rules to avoid their effect (one regressor for each of two colours from different rules). For the RSA GLMs, each trial contributed to the estimation of 2 regressors (e.g., coarse stimulus information and rule). In all GLMs, movement parameters, block means, and run means were also included as covariates of no interest, and trials were modelled as epochs lasting from stimulus onset until response (Grinband et al., 2008). Error trials were excluded from the analysis. We estimated the regressors for each of the 12 blocks of each participant obtaining 12 beta values which we used in MVPA.

### Analysis 1: Multivariate pattern analysis (MVPA)

We used a standard multivariate pattern decoding to quantify the level of different types of information consisting of coarse and fine stimulus, rule and response information, as defined above, in fMRI (Figure 2A) and MEG (Figure 2B).

In fMRI, we used a leave-one-block-out cross-validation approach for decoding using a linear SVM classifier as implemented in Matlab. This way, we trained the classifier on all-minus-one blocks and tested the classifier on the remaining one block and repeated the procedure until all blocks were used in testing once. The decoding accuracy for a single participant was calculated by averaging the decoding results across cross-validation runs. The number of voxels within each ROI determined the dimension of the data entered into the classifier. The classification was performed for each ROI in each hemisphere separately and then averaged across the two hemispheres and reported.

In MEG, we used a time-resolved decoding procedure (Robinson et al., 2022), where we repeated the decoding analysis on every time point along the trial, once using the data aligned to the stimulus onset time (*stimulus-aligned* analysis) and once with the data aligned to the response time of each trial (*response-aligned* analysis). The two different alignments provide insights about the temporal profile of information encoding affected by stimulus onset and those aligned to the behavioural outputs. We used a 10-fold cross-validation procedure, with 9 folds of the data used for training the classifier and the left-out fold used for testing it and repeating the procedure 10 times until all folds are used once in testing. The decoding accuracy for a single participant was calculated by averaging the decoding results across these cross-validation runs.

### Analysis 2: RSA-based fMRI-MEG fusion

We used RSA to provide a common platform for comparing the representation of information across ROIs in fMRI and across time in MEG. For each participant (different in fMRI and MEG), ROI in fMRI, or time point in MEG, we constructed four distinct neural representational dissimilarity matrices (RDMs) by decoding different combinations of stimuli, cues, rules and responses (Figure 3A). These 4 RDMs were designed to capture coarse and fine stimulus, rule and response information. Each cell in the RDM reflects the decoding accuracy of the two conditions of the column and row of the matrix. For example, the decoding value at the intersection of column “Right stim. Rule 1” and row “Left stim. Rule 1” contains the decoding of right stimuli in rule 1 and left stimuli in rule 1, therefore reflecting information (dissimilarity) about stimulus side irrespective of other variables such as rule. We obtained fMRI neural RDMs for every ROI and then averaged them across hemispheres. We also obtained MEG neural RDMs for every time point across the trial, once for the *stimulus-aligned* data and once for *response-aligned* data. MEG RDMs were constructed using signals from the sensors over the whole brain.

We also constructed 4 theoretical model RDMs to quantify coarse and fine stimulus, rule and response information within neural RDMs explained above (Figure 3A). For example, in the *coarse stimulus information* model RDM, the elements which corresponded to the decoding of right stimuli from right stimuli, or left stimuli from left stimuli representations (and not their cross conditions) were valued as 0.5, and the elements which corresponded to the cross-conditions between right and left stimuli were valued as 1. For RSA, fusion and connectivity analyses (explained below), we selected and reshaped the lower triangular elements of the RDMs (excluding the diagonal elements) into vector RDMs (RDVs).

To fuse fMRI and MEG representations, we first averaged the RDM across participants and used a model-based approach using the subject-average neural fMRI and MEG RDVs and the model RDVs (Hebart et al., 2017, Moerel et al., 2024). To that end, we used *Spearman’s* partial correlation analysis and obtained one commonality index for each ROI in fMRI at each time point of MEG and using each of the four models as defined in (1):

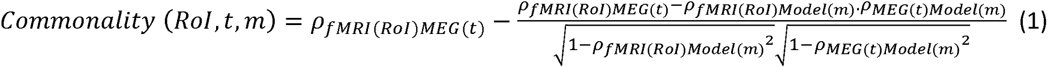

where *ρ_fMRI(RoI)MEG(t)_* refers to the Spearman correlation between the neural fMRI RDV in region *RoI* and neural MEG RDV at time *t*, *ρ_fMRI(RoI)Model_*_(*m*)_ refers to the neural fMRI RDV in region *RoI* and model RDV *m*, and *ρ_MEG(t)Model_*_(*m*)_ refers to the neural MEG RDV at time *t* and model RDV *m* . Using this formula, in which the fraction term is simply the correlation between the fMRI and MEG RDVs where model RDV is partialled out, we obtain four commonality indices for the four models in each ROI at every 5 ms time point. Equation 1 results in the brown area indicated in the Venn diagram shown in Figure 4A.

### Analysis 3: Fusion-based representational connectivity analysis (Fusion-RCA)

We developed the Fusion-RCA to investigate how representation/information is transferred within the MDN and between the MDN and visual areas. While several recent RCA approaches have been used for the evaluation of connectivity, they all used one brain imaging modality. Specifically, while the concept of RCA is not specific to the imaging modality and can potentially be applied to all multivariate recording modalities, all previous methods used either fMRI (Coutanche et al., 2020) or E/MEG (Goddard et al., 2016; Karimi-Rouzbahani, 2018; Karimi-Rouzbahani et al., 2019; Kietzmann et al., 2019; Karimi-Rouzbahani et al., 2021a; Karimi-Rouzbahani et al., 2021b; Goddard et al., 2022; Karimi-Rouzbahani et al., 2022), which respectively lack the temporal and spatial resolution needed to track rapid and spatially fine-grained cognitive processes in the brain.

Here we utilise the spatiotemporal specificity of the commonality time courses, which we obtain through fMRI-MEG fusion, and the logic of Granger causality, to track information across the brain with more spatial and temporal resolution than is achievable from either fMRI or MEG alone. Specifically, to evaluate information flow, we evaluated Granger causal relationships between the time courses of commonality indices from different ROIs and for different types of information separately (Figure 4A). We use the multivariate Granger causality (MVGC) toolbox (Barnett and Seth, 2014) which allows the evaluation of Granger causality between multiple areas simultaneously. Granger causality suggests that if signal (here commonality index) X at the present time point improves the prediction of future time point of signal Y beyond present time point of signal Y, then X is said to Granger cause Y. As an advantage over conventional implementations of Granger causality, which involve the estimation of regression parameters once for the full model and once for the reduced model, MVGC does not explicitly estimate the reduced model, eliminating one source of estimation error leading to improved statistical power. It is achieved through the calculation of multiple equivalent representations of vector autoregressive models (Barnett and Seth, 2014), which is the core of Granger causality prediction. Application of MVGC to the commonality indices results in matrices of connectivity (Figure 4A), reflecting measured Granger causal relationship/connectivity between source and destination ROIs, and determining the direction of causality/information flow. Note that as the commonality indices used in our new connectivity analysis are already specific about the type of information they reflect, the resultant connectivity quantifies the potential flow of information, rather than activations as conventionally measured by connectivity measures. For a review of caveats and nuances of model-based and model-free RCA see Karimi-Rouzbahani et al. (2022). After statistical analysis (below) we visualised the connectivity results into an Information Transfer Diagram in which significant Granger-causal influences are depicted with an arrow (Figure 4C).

### Statistical analyses

#### Bayes factor analysis

We used Bayes factor t-tests, as implemented by Bart Krekelberg^2^ based on Rouder et al. (2012), for comparing the levels of decoding across conditions and between a condition and chance decoding. We used standard rules of thumb for interpreting levels of evidence (Lee and Wagenmakers, 2005; Dienes, 2014): Bayes factors of >3 and <1/3 were interpreted as evidence for the alternative and null hypotheses, respectively. We interpreted Bayes factors between 3 and 1/3 as insufficient evidence either way.

To evaluate the evidence for the null and alternative hypotheses of at-chance and above-chance decoding in fMRI, respectively, we generated a null distribution containing 1000 decoding values obtained by randomising class labels 1000 times (random permutation). As we were also interested in evaluating the effect of perceptual difficulty on information coding, we also used Bayes factor analysis to evaluate the evidence for difference in decoding levels between coarse and fine stimulus information across participants in each ROI separately.

To evaluate the evidence for the null and alternative hypotheses of at-chance and above-chance decoding in MEG, respectively, we generated a null distribution containing 1000 decoding values obtained by randomising class labels 1000 times (random permutation) for every time point. As in fMRI, to evaluate the evidence for difference in decoding levels between coarse and fine stimulus information, we compared their decoding rates across participants at every time point. Accordingly, we performed the Bayes factor analysis for alternative (i.e., difference; H1) versus the null (i.e., no difference; H0) hypotheses.

The priors for all Bayes factor analyses were determined based on Jeffrey-Zellner-Siow priors (Jeffreys, 1998; Zellner and Siow, 1980) from the Cauchy distribution based on the effect size that is initially calculated in the algorithm using t-test (Rouder et al., 2012). The priors are data-driven and have been shown to be invariant with respect to linear transformations of measurement units (Rouder et al., 2012), which reduces the chance of being biased towards the null or alternative hypotheses. We did not perform correction for multiple comparisons when using Bayes factors as they are much more conservative than frequentist analysis (Gelman and Tuerlinckx, 2000; Gelman et al., 2012).

#### Permutation testing for fusion

To evaluate the significance of commonality indices obtained using partial correlation, we tested the true partial correlations against a null distribution obtained by shuffling the class labels in fMRI data and regenerating the RDMs using the data with shuffled labels 1000 times. We compared the true commonality indices at every time point with the randomly generated commonality indices for the same time point and deemed it significant if it exceeded 95% of the random correlations (p < 0.05) after correcting for multiple comparisons across time (using Matlab mafdr function which uses the direct approach of Storey, 2002, where the algorithm fixes the rejection region and then estimates its corresponding error rate resulting in increased accuracy and power). We used a permutation approach rather than Bayes factor analysis here as there was no chance level of commonality known *a priori*. For quantitative comparison of commonality indices, we extracted three parameters from the commonality time courses consisting of “time to first significant commonality” (time from stimulus onset to the first significant commonality), “time to maximum commonality” (time from stimulus onset to the maximum commonality) and “time from maximum commonality” (time from the maximum commonality to the response).

#### Connectivity statistics

For statistical testing of connectivity results, we used the statistical test results produced by the MVGC toolbox. The significance of connectivity indices obtained from the MVGC is evaluated based on a theoretical (asymptotic) null distribution which is corrected for multiple comparisons based on the number of source and destination ROIs, number of time points within the trial, the number of trials and the order of the auto-regression model (Barnett and Seth, 2014). The method shuffles the time samples in the time series (commonality indices here) to generate null time series from the true time series and evaluates the significance of the true connectivity values against those obtained from the null time series.

## Acknowledgements

The authors thank Dr Amanda Robinson for providing the MEG data and the experimental scripts and for helpful discussion of the experimental design. This work was supported by Australian Research Council (ARC) Discovery Project grant DP170101840 awarded to AW and ANR. HK-R was further supported by Newton International Fellowship (NIF\R1\192608) and follow-on funding (AL\231037) from the Royal Society. AW was supported by Medical Research Council (UK) intramural funding SUAG/093/G116768 and an ARC Future Fellowship FT170100105. ANR is supported by an ARC Future Fellowship FT230100119. The authors acknowledge the facilities and scientific and technical assistance of the National Imaging Facility, a National Collaborative Research Infrastructure Strategy (NCRIS) capability, at Macquarie Medical Imaging and the KIT-Macquarie Brain Research Laboratory, Macquarie University. For the purpose of open access, the authors have applied a Creative Commons Attribution (CC BY) licence to any Author Accepted Manuscript version arising from this submission.

## Footnotes

1 Information which is represented more strongly can suppress weaker information when the former is partialled out from the latter in RSA, making it harder to detect the weaker information.

2 https://klabhub.github.io/bayesFactor/

## Notes

### Competing Interest Statement

The authors have declared no competing interest.

